# DNA-PKcs is required for cGAS/STING-dependent viral DNA sensing in human cells

**DOI:** 10.1101/2022.11.01.514678

**Authors:** Dayana Hristova, Marisa Oliveira, Emma Wagner, Alan Melcher, Kevin Harrington, Alexandre Belot, Brian J Ferguson

**Affiliations:** Department of Pathology, University of Cambridge, Tennis Court Road, Cambridge, CB2 1QP, UK; The Institute of Cancer Research, London SW7 3RP, UK; Centre International de Recherche en Infectiologie, Inserm, U1111, Université Claude Bernard, Lyon, France

**Keywords:** DNA, innate immunity, virus, DNA-PK, STING, cGAS

## Abstract

To mount an efficient interferon response to virus infection, intracellular pattern recognition receptors (PRRs) sense viral nucleic acids and activate anti-viral gene transcription. The mechanisms by which intracellular DNA and DNA viruses are sensed are relevant not only to antiviral innate immunity, but also to autoinflammation and anti-tumour immunity through initiation of sterile inflammation by self-DNA recognition. The PRRs that directly sense and respond to viral or damaged self-DNA function by signalling to activate interferon regulatory factor (IRF)-dependent type one interferon (IFN-I) transcription. We and others have previously defined DNA-dependent protein kinase (DNA-PK) as an essential component of the DNA-dependent antiviral innate immune system. Here, we show that DNA-PK is essential for STING-dependent IFN-I responses in human cells during stimulation with exogenous DNA and infection with DNA viruses.

## Introduction

The ability of cells to sense and respond to pathogens by producing type I interferons (IFN-I) and inflammatory mediators is essential for host defence against infection. Pattern recognition receptors (PRRs) that bind nucleic acids and drive the transcription of IFN-I are specifically required to control virus infections^1^. IFN-I acts in an autocrine or paracrine fashion to upregulate the expression of proteins that establish the anti-viral state in infected tissues, whilst the secretion of chemokines and cytokines attracts and activates tissue-resident and circulating leukocytes to help clear the infection^2^.

In the initial events of detection of viruses, genomic nucleic acids trigger the activation of PRRs that bind DNA or RNA directly and signal downstream to activate the transcription factors interferon regulatory 3 (IRF3) and nuclear factor kappa B (NF-κB). These active transcription factors move to the nucleus to initiate interferon and inflammatory gene activation. In the context of DNA virus infections, multiple receptors that activate this pathway have been identified. The DNA-binding PRRs cyclic GMP-AMP synthase (cGAS), DNA-PK and interferon-inducible protein 16 (IFI16) function to sense and respond to DNA-containing pathogens and drive the innate and subsequent adaptive immune response to virus infection^3–9^.

The DNA-dependent protein kinase (DNA-PK) complex consists of a phosphatidylinositol 3-kinase-related protein kinase (PIKK) family catalytic subunit called DNA-PKcs, and two DNA-binding regulatory subunits Ku70 and Ku80. This heterotrimeric complex functions in non-homologous end joining (NHEJ) that repairs nuclear double-stranded DNA breaks by directly binding broken ends of genomic DNA. We previously defined a function for DNA-PK in the sensing of intracellular DNA and DNA virus infections via activation of an IRF3-dependent pathway^4^. Others have re-affirmed the function of DNA-PK in sensing viruses and established DNA-PK as a critical regulator of antiviral immunity^7–12^. In order to activate IRF3, viral DNA-sensing PRRs signal via activation of an adaptor protein, stimulator of interferon genes (STING)^13^. STING binds the second messenger 2’-3’cGAMP, a product of the enzyme activity of cGAS^14^ that subsequently drives downstream IRF3 and NF-κB activation by a mechanism that requires the IKK family kinases including TANK-binding kinase-1 (TBK-1)^15,16^. DNA-PK has been reported to act in STING-dependent and -independent pathways depending on the cell type and activation context^17,18^, and to activate IRF3 but not NF-κB-dependent signalling during infection or stimulation^4^. DNA-PK is also itself a target of multiple viral counter-defence mechanisms encoded by large DNA viruses. Poxviruses and herpesviruses encode proteins that bind and/or target DNA-PK for degradation during infection^19–21^, providing further evidence for the importance of this complex in regulation of anti-viral immunity.

Here, we establish that DNA-PK functions in human cells to sense and respond to intracellular dsDNA and to the vaccinia virus (VACV) and herpes simplex virus 1 (HSV-1) viral DNA to drive interferon production. DNA-PK functions in the same pathway as cGAS and STING in this context and is required for STING activation and subsequent IRF3-dependent interferon and chemokine production. This work thereby establishes the function of DNA-PK in STING-dependent anti-viral immune responses in human cells.

## Methods

### Cells

Human immortalised foreskin fibroblasts (HFF-hTert), U20S, BSC-1, baby hamster kidney (BHK) and primary skin fibroblasts were cultured in DMEM with 10% v/v foetal calf serum (FCS) and 50 μg/mL Pen-strep. Chicken embryonic fibroblasts (CEF) were cultured in DMEM-F12 with Glutamax (Gibco), 5% v/v FBS, and 50 μg/mL pen-strep. Fibroblast from patients carrying biallelic p.Leu3062Arg mutation were provided by the CRB Biotech, Lyon, France. The study was approved by the Medical Ethics Committee of Sud Est III (Lyon, France) and carried out in accordance with the Declaration of Helsinki principles. Patient provided written informed consent for inclusion of their details and samples in the study.

### Viruses

Modified vaccinia Ankara (MVA) was grown on BHK cells and titrated on primary CEFs. Titrations were counted by immunostaining using an anti-VACV Lister cocktail antibody (RayBiotech, MD-14-1041), secondary anti-rabbit horseradish peroxidase (HRP)-conjugated antibody (Sigma, A6154) and True-blue substrate (KPL) to visualise plaques. VACV strain TBio 6517, currently in early clinical testing (NCT04301011), kindly provided by Turnstone Biologics, was grown and titrated on BSC-1 cells. For growth curve analysis, HFF-hTERT cells were infected with VACV TBio 6517 at multiplicity of infection (MOI) 0.01. 24 or 48 hours later, cell lysates were prepared, frozen and thawed three times and sonicated to obtain cell-associated virus. Numbers of infectious virions were quantified by titration on BSC-1 cells. HSV-1 strains S17 (wild type) and *dl403* lacking the *ICP0* gene (HSV-1ΔICP0)^22^ were grown and titrated on U2OS cells and plaques were counted using toluidine blue staining.

### Knockout cell line generation

HFF-hTERT-Cas9 cells were transduced with lentiviruses expressing gRNAs targeting *PRKDC*, *TMEM173* or *MB21D1* (Table 1). Cell lines were selected with puromycin and analysed by immunoblotting for successful editing.

**Table 1:**
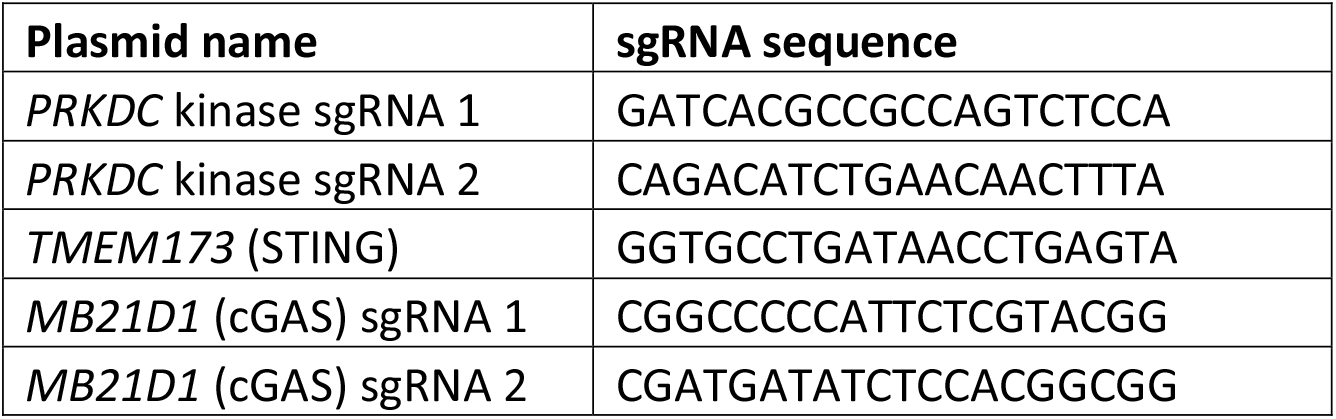
sgRNA primer sequences.

### Stimulations

HFF were seeded in tissue culture plates and, the following day, cells were transfected using TransIT-LT1 (Mirus Bio, USA) with herring testis (HT)-DNA, calf thymus (CT)-DNA, or Poly(I:C) and harvested 6 or 16 h post-transfection.

### qRT-PCR

Cells were lysed in 250 μL of lysis buffer containing 4 M guanidine thiocyanate, 25 mM Tris pH 7, and 143 mM 2-mercaptoethanol and purified on silica columns (Epoch Life Science). Using 500 ng of RNA, cDNA was produced using SuperScript III reverse transcriptase, following the manufacturer’s protocol (Thermo Scientific, Waltham, MA, USA). cDNA was used for quantitative PCR (qPCR) in a final volume of 10 μL. qPCR was performed using SybrGreen Hi-Rox (PCR Biosystems Inc.) using primers described in Table 2. Fold change in mRNA expression was calculated by relative quantification using GAPDH as endogenous control.

**Table 2:**
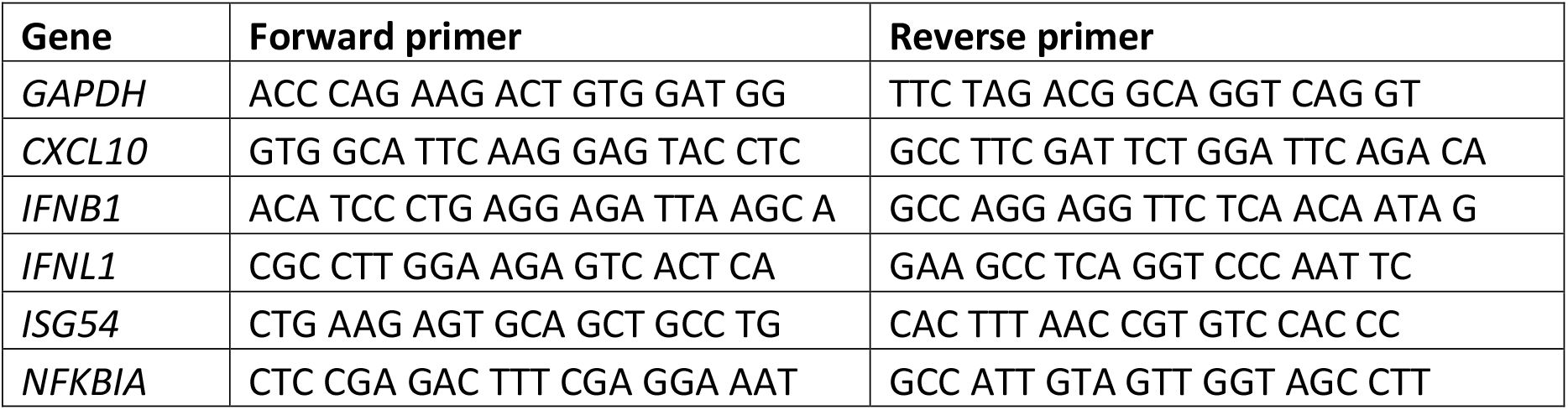
qRT-PCR primer sequences.

### Immunoblotting

Cells were lysed in radioimmunoprecipitation assay (RIPA) lysis buffer (50 mM Tris-HCL pH8, 150mM NaCl, 1% NP-40, 0.1% SDS, 0.5% Na Deoxycholate), cOmplete Mini EDTA-free protease inhibitors (Roche), as well as PhosSTOP phosphatase inhibitor cocktail (Roche) and quantified using the bicinchoninic (BCA) assay (Thermo Scientific). Protein samples were run on 10-12% SDS polyacrylamide gels on a Bio-Rad Protean III system or 4-12% bis-Tris gradient Nu-PAGE gels on a Novex Mini-Cell system (Invitrogen) (Invitrogen). After electrophoresis, proteins were transferred onto nitrocellulose membrane, immunoblotted with the indicated antibodies (Table 3) and imaged by a Li-Cor Odyssey CLx.

**Table 3:** Antibodies used in this study.

| **Antibody target** | **Source and species** |
| --- | --- |
| cGAS | Santa Cruz (sc-515777); mouse |
| DNA-PKcs (cocktail) | Thermo Scientific (ms-423-p1); mouse |
| IFI16 | Santa Cruz (sc-8023); mouse |
| IκBα | Santa Cruz (sc-1648); mouse |
| IRF3 | Abcam (ab68481); rabbit |
| Ku70 | Abcam (ab3114); mouse |
| TBK1 | Abcam (ab40676); rabbit |
| STING | Cell Signalling (13647); rabbit |
| Tubulin | Millipore (05-829); mouse |
| HSV-1 ICP0 | Santa Cruz (sc-53070); mouse |
| VACV Lister | RayBiotech (MD-14-1041); rabbit |
| p-IRF3 (Ser386) | Abcam (ab76493); rabbit |
| p-TBK1 (Ser172) | Cell Signalling (5483S); rabbit |
| p-STING (Ser366) | Cell Signalling (85735); rabbit |
| γH2AX | Millipore (05-636); mouse |
| Anti-mouse IgG antibody (800) | Li-Cor (926-32210); goat |
| Anti-rabbit IgG antibody (680) | Li-Cor (926-68071); goat |
| Anti-mouse IgG | Alexa Fluor 488 (A21202); donkey |
| Anti-mouse IgG | Alexa Fluor 546 (A10036); donkey |
| Anti-rabbit IgG | Alexa Fluor 546 (A10040); donkey |

### Immunofluorescence microscopy

Cells were seeded in a 24-well plate on sterile 13 mm coverslips and fixed for 10 minutes with cold 4% w/v formaldehyde (Fisher Scientific) and cold 8% w/v formaldehyde in HEPES buffer. Cells were permeabilised for 5-10 minutes with 0.25% v/v Triton X-100 in PBS, blocked with 5% w/v milk in PBS at RT for one hour. The cells were then incubated overnight at 4°C with primary antibody (Table 3) at the indicated dilution in 1% w/v milk in PBS. Following this, the cells were washed with PBS three times and were incubated at RT for 30 minutes in the dark with secondary antibody diluted 1:1000 in 1% w/v milk in PBS. Coverslips were mounted onto slides with 10 μL of mounting solution (25 % glycerol v/v, 0.1 M Tris pH 8.5, 10 % Mowiol 4-88 w/v containing 4’, 6-diamidino-2-phenylindole (DAPI). A Zeiss Pascal Confocal microscope was used to visualise the samples.

## Results

### Human fibroblasts respond to intracellular DNA (ICD) by activating STING/TBK1/IRF3

To establish a model system for studying the innate immune function of DNA-PK in human cells, we assessed multiple human cell lines for their ability to activate an interferon response to intracellular dsDNA. We found that, although many human cell lines did not respond to transfection of dsDNAs, human foreskin fibroblasts (HFF) mounted a robust response. Fibroblasts are primary targets of virus infection and help coordinate multiple aspects of innate immunity and inflammation^4,23,24^. Following dsDNA transfection, HFFs activate STING, TBK1 and IRF3, as evidenced by phosphorylation of these signalling pathway components (Figure 1A). Activation of this pathway leads to IFN-I and chemokine (CXCL10) transcription (Figure 1B), key target genes directly transcriptionally activated by ICD-dependent IRF3 activity^4^, although we did not detect any induction of type III interferon or interferon alpha above background levels, and only very low levels of NF-κB-dependent transcripts, such as *NFKBIA* (Figure 1B).

**Figure 1.**
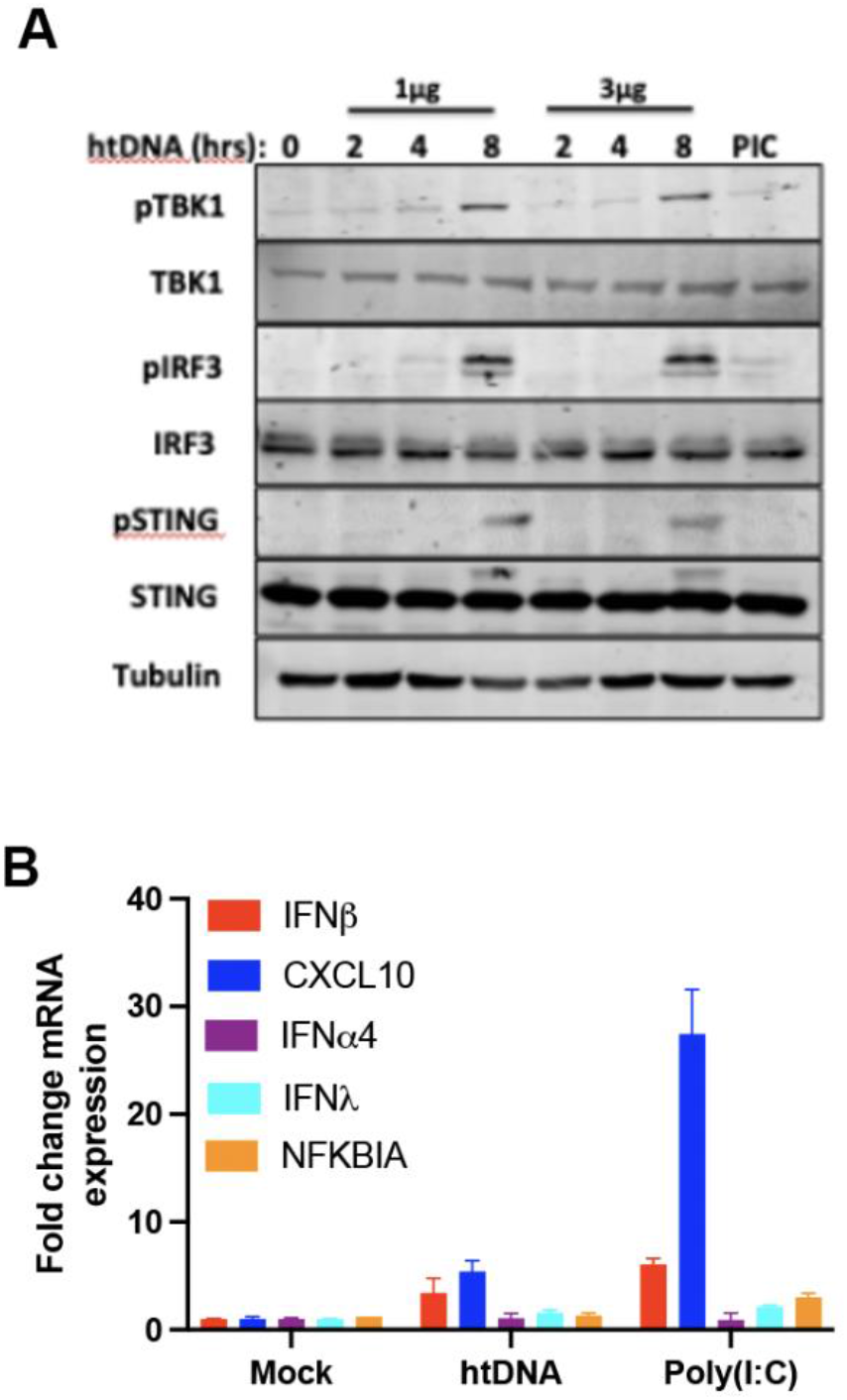
Human fibroblasts respond to ICD stimulation by activating the STING/IRF3/TBK1 signalling axis. A) HFFs were stimulated by transfection with 1 or 3 μg/ml of HT-DNA for the indicated times and immunoblotted for the indicated proteins. B) HFFs were stimulated by transfection with the indicated DNAs or poly(I:C) and analysed by qRT-PCR 6 h later for the indicated genes. n=3 *p<0.5, **p<0.01.

### ICD sensing in human fibroblasts requires cGAS, STING and TBK1/IKKe

We next set out to define the pathway leading to IRF3 activation in response to ICD-induced stimulation in HFFs. To do this, we created STING and cGAS knockout lines and stimulated them by transfection with HT-DNA. cGAS was required for ICD-driven interferon and chemokine transcription (Figure 2A). Analysis of IRF3 activation status showed that, while ICD stimulation of cGAS KO HFFs showed little or no IRF3 activation, the ability of these cells to respond to RNA was maintained (Figure 2B). STING KO cells also lost ICD-driven interferon responses (Figure 2C) and use of the TBK1/IKK*ε* inhibitor, BX795, indicated that this ICD sensing pathway is also completely dependent on these kinases (Figure 2D). As such, we show here that the ICD pathway in human fibroblasts uses the canonical cGAS/STING/TBK1/IRF3 signalling axis.

**Figure 2.**
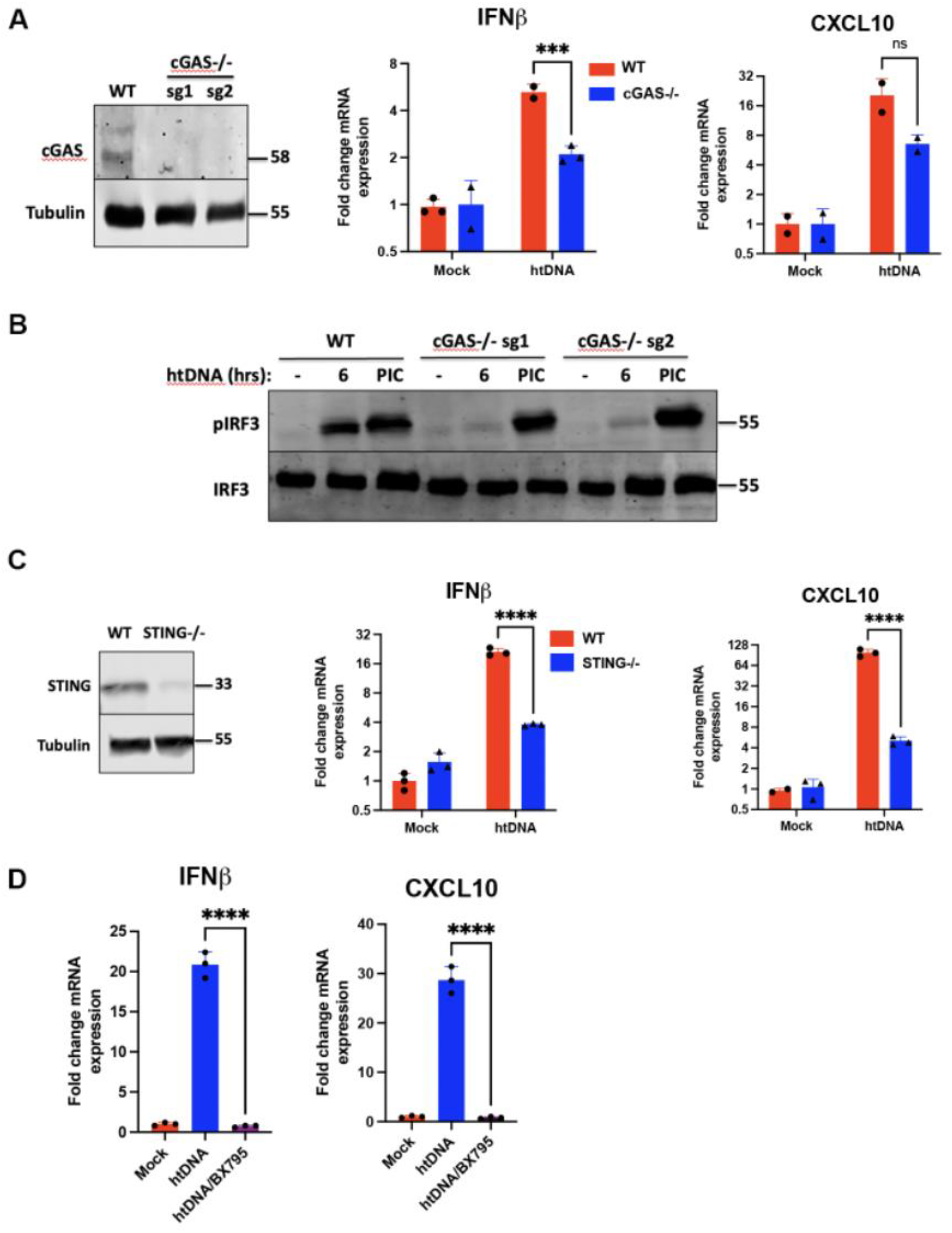
The interferon response to ICD stimulation in HFFs is dependent on cGAS, STING and TBK1. A) cGAS knockout cell lines were generated by CRISPR/Cas9 with two guide RNAs targeting different exons of the *MB21D1* gene (sg1 and sg2) and were immunoblotted with anti-cGAS antibody. WT (cGAS^+/+^) and cGAS^−/-^ cells were stimulated with HT-DNA and analysed by qRT-PCR 6 h later for transcription of *IFNB* and *CXCL10*. n=3 ***p<0.05. B) WT (cGAS^+/+^) and cGAS^−/-^ cells were stimulated with HT-DNA or poly(I:C) and analysed by immunoblotting with the indicated antibodies. C) STING knockout cells were generated by CRISPR/Cas9 with a guide RNA of the *TMEM173* gene and were immunoblotted with an anti-STING antibody. WT (STING^+/+^) and STING^−/-^ cells were stimulated with HT-DNA and analysed by qRT-PCR 6 h later for transcription of *IFNB* and *CXCL10*. n=3 ****p<0.005. D) WT HFFs were pre-treated with the TBK1 inhibitor BX795, stimulated by transfection with HT-DNA and analysed by qRT-PCR 6 h later for transcription of *IFNB* and *CXCL10*.

### DNA-PKcs is required for STING activation and IFN-I production in response to ICD stimulation

To assess the role of DNA-PK in DNA sensing in human cells, we stimulated HFFs with ICD and monitored activation of DNA-PKcs using an autophosphorylation-specific antibody that recognises pS2056. Stimulation of cells with ICD or using etoposide to initiate DNA damage resulted in an increase in pS2056 signal above background levels (Figure 3A) indicating activation of DNA-PK, a process that usually occurs via DNA end binding. We next generated two CRISPR KO cells lines using sgRNAs targeting different DNA-PKcs exons (Figure 3B). Stimulation of these DNA-PKcs KO human fibroblasts resulted in complete loss of *IFNB* and *CXCL10* transcription (Figure 3C) and, also, reduction in transcription of the IRF3-specific target gene *ISG54* (Figure 3C). Analysis of intracellular signalling pathway activation in these cells indicated that DNA-PKcs loss resulted in significant reductions in ICD-driven STING and IRF3 activation, although intracellular RNA-driven IRF3 activation was unaffected (Figure 3D). As such, these data indicate that DNA-PKcs is essential for the cGAS/STING pathway of ICD sensing in human cells.

**Figure 3.**
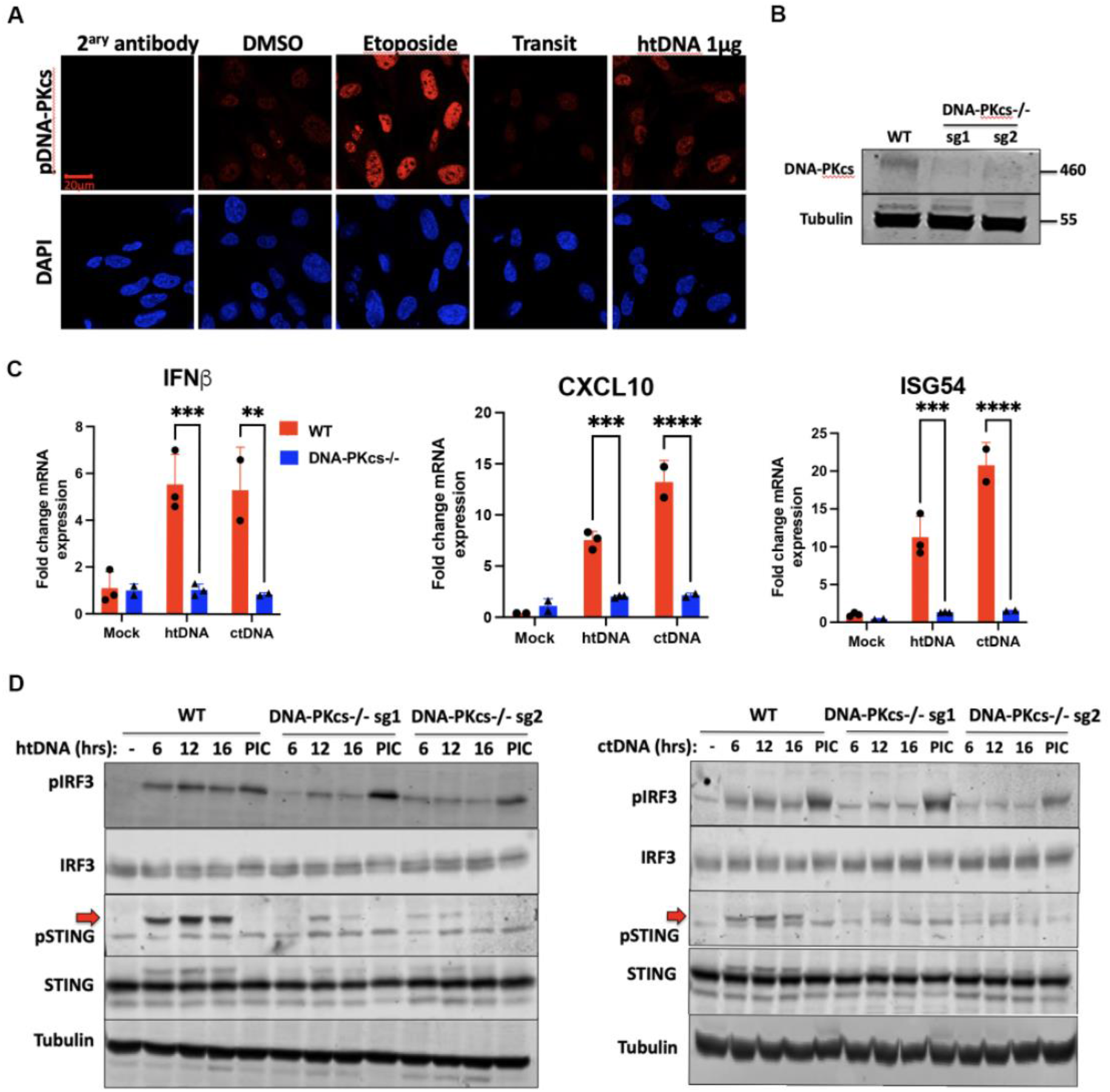
DNA-PKcs is required for STING-dependent sensing in human fibroblasts. A) DNA-PKcs is activated during ICD stimulation. Representative images of cells treated with etoposide, HT-DNA or carrier controls, fixed and stained by immunofluorescence with an antibody recognising phospho-serine 2056 on DNA-PKcs. Cells are counterstained with DAPI, scale bar = 20 μm. B) DNA-PKcs knockout cell lines were generated by CRISPR/Cas9 with two guide RNAs targeting different exons of the *PRKDC* gene (sg1 and sg2) and were immunoblotted with anti-DNA-PKcs antibody. C) WT (DNA-PKcs^+/+^) and DNA-PKcs^−/-^ cells were stimulated with HT-DNA or CT-DNA and analysed by qRT-PCR 6 h later for transcription of *IFNB*, *ISG54* and *CXCL10*. n=3 **p<0.01, ***p<0.05, ****p<0.01. D) WT (DNA-PKcs^+/+^) and DNA-PKcs^−/-^ cells were stimulated with ctDNA (top panel) or HT-DNA (bottom panel) or Poly(I:C) (PIC) for the indicated times and analysed by immunoblotting with the indicated antibodies. Red arrows show the position of the phospho-STING specific band.

The kinase activity of DNA-PKcs is required for multiple aspects of DNA-PK function, including its roles in DNA damage repair and in V(D)J recombination. Here, we asked whether inhibition of the DNA-PKcs kinase domain with small molecule inhibitors could impact its function in ICD sensing. We first used the inhibitor NU7441 that can inhibit DNA-PKcs activity (Supplementary Figure 1A) and found that pre-treatment of cells with NU7441 enhanced the ICD-driven interferon responses (Supplementary Figure 1B). As this result was incongruous with our data from DNA-PKcs KO cells, we hypothesised that this compound may have an off-target effect that impacts this signalling pathway. Indeed, we found that NU7441 could activate TBK1 in the absence of DNA-PKcs (Supplementary Figure 1C), indicating a strong off-target effect on the ICD pathway of this molecule and, hence, it is unsuitable in this context. As an alternative, we used the more-specific compound AZD7648^25^ that can inhibit DNA damage-dependent histone phosphorylation (γH2AX) (Supplementary Figure 2), a process that is DNA-PKcs-dependent, but had no impact on ICD-dependent TBK1 and IRF3 phosphorylation (Figure 4A), or *IFNB* transcription (Figure 4B) in HFFs. As such, the kinase activity of DNA-PKcs was not found to be required in the context of DNA sensing in human fibroblasts.

**Figure 4.**
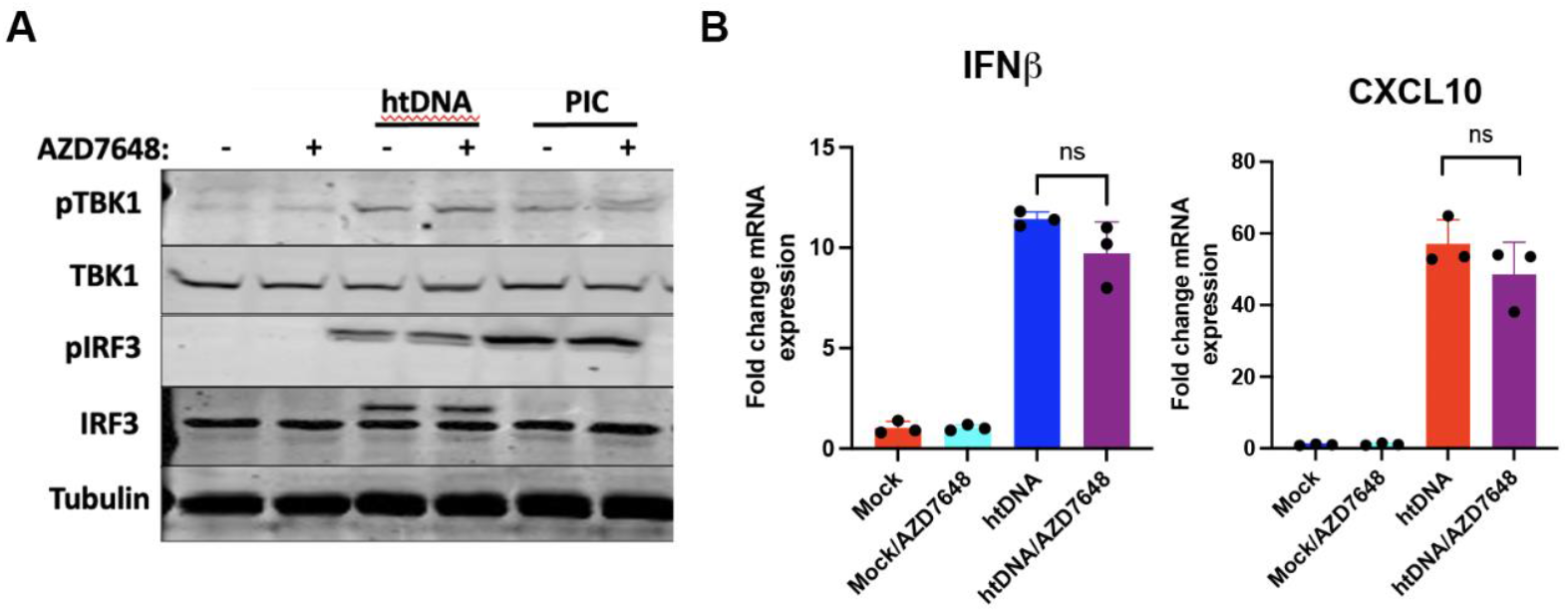
DNA-PKcs kinase activity is dispensable for ICD sensing. Cells were pre-treated with AZD7648 for 1 hour prior to transfection with HT-DNA or PIC and A) immunoblotted for the indicated antibodies or B) analysed by qRT-PCR for *IFNB* and *CXCL10* 6 h post stimulation.

### The human mutation DNA-PKcs L3062R is a gain of function mutation for ICD sensing

Loss of DNA-PKcs function is associated with severe-combined immunodeficiency disorder (SCID), characterised by loss of B and T cell repertoires due to a lack of V(D)J recombination. Since the discovery that DNA-PKcs is essential for V(D)J recombination in mice, several human polymorphisms have been discovered that result in a similar primary immune deficiency. Once such rare mutation is L3062R^26^. This polymorphism results in human SCID and is associated with a loss of the DNA repair function of DNA-PKcs, resulting in failed V(D)J recombination. The amino-acid residue L3062 lies on the surface of DNA-PKcs in a region of the protein associated with Artemis-binding and does not impact the kinase activity. Of interest is the observation that patients carrying the biallelic L3062R DNA-PKcs variant exhibit interferon signatures in the blood^27^. Interestingly, this positive signature persists three years after the bone marrow transplantation in one patient (IFN signature > 16, normal value <2.3). We analysed the ability of primary skin fibroblasts from healthy donor or a patient carrying this mutation for their response to ICD stimulation. Cells carrying the L3062R mutation showed enhanced STING/TBK1/IRF3 activation and *IFNB* transcription in response to ICD stimulation (Figure 5A, B). Cells carrying this variant also showed enhanced CXCL10 transcription when resting, indicating increased basal inflammatory signalling (Figure 5B). These data provide further direct evidence of the function of DNA-PKcs in regulation of ICD sensing in humans.

**Figure 5.**
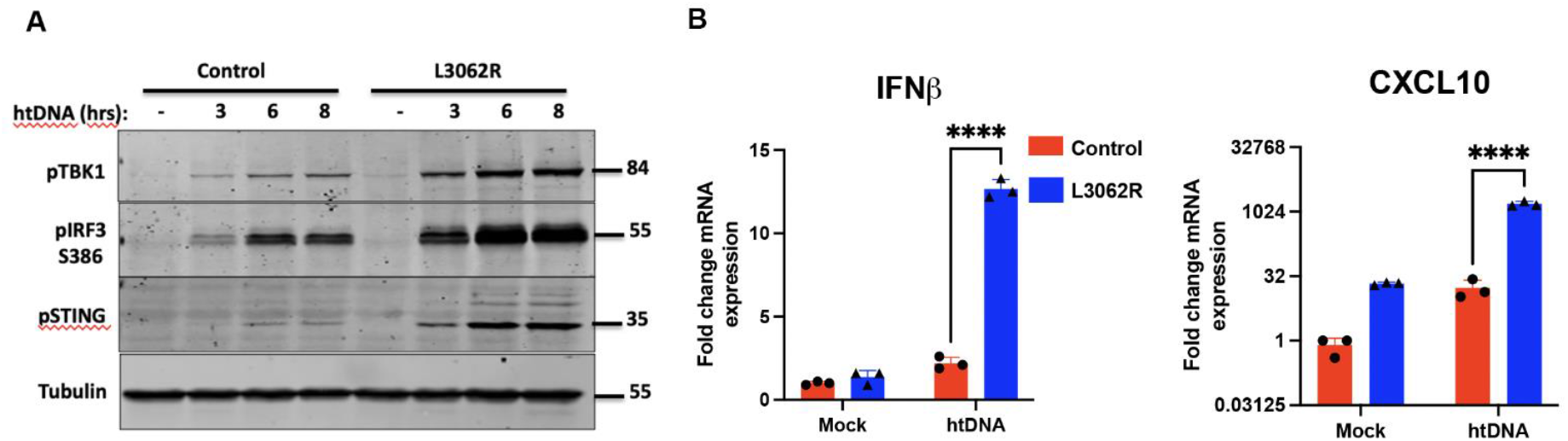
The human DNA-PKcs L3062 mutation enhances STING signalling. A) Primary skin fibroblasts from healthy donor or a patient harbouring the L3062 mutation were stimulated with HT-DNA and A) immunoblotted with the indicated antibodies or B) analysed by qRT-PCR for *IFNB* and *CXCL10* 6 h post stimulation.

### DNA-PKcs is essential for STING-dependent sensing of DNA viruses

To determine the contribution of DNA-PKcs to the innate sensing of DNA viruses, we used two infection models with dsDNA viruses. Modified vaccinia Ankara (MVA) is a derivative of the vaccinia virus strain chorioallantois vaccinia Ankara (CVA) produced by extensive passage in chicken cells. MVA has lost numerous immunomodulatory proteins as well as the ability to replicate in human cells. MVA can still enter most human cells, however, and presents large amounts of viral DNA into the cytoplasm that can activate cGAS/STING signalling^4,28,29^. Infection of HFFs with MVA resulted in STING and IRF3 phosphorylation that was markedly reduced in the absence of DNA-PKcs (Figure 6A), indicating that DNA-PKcs is required for vaccinia virus-driven activation of the STING/TBK1/IRF3 signalling axis. As MVA is non-replicative in human cells, we used a replicating strain of VACV (TBio 6517) to analyse the impact of DNA-PKcs loss on the yield of virus from infected cells. In a multi-step growth curve analysis, we found that cells lacking DNA-PKcs produced a significantly enhanced virus yield (Figure 6B), consistent with its role in host defence against DNA virus infections.

**Figure 6.**
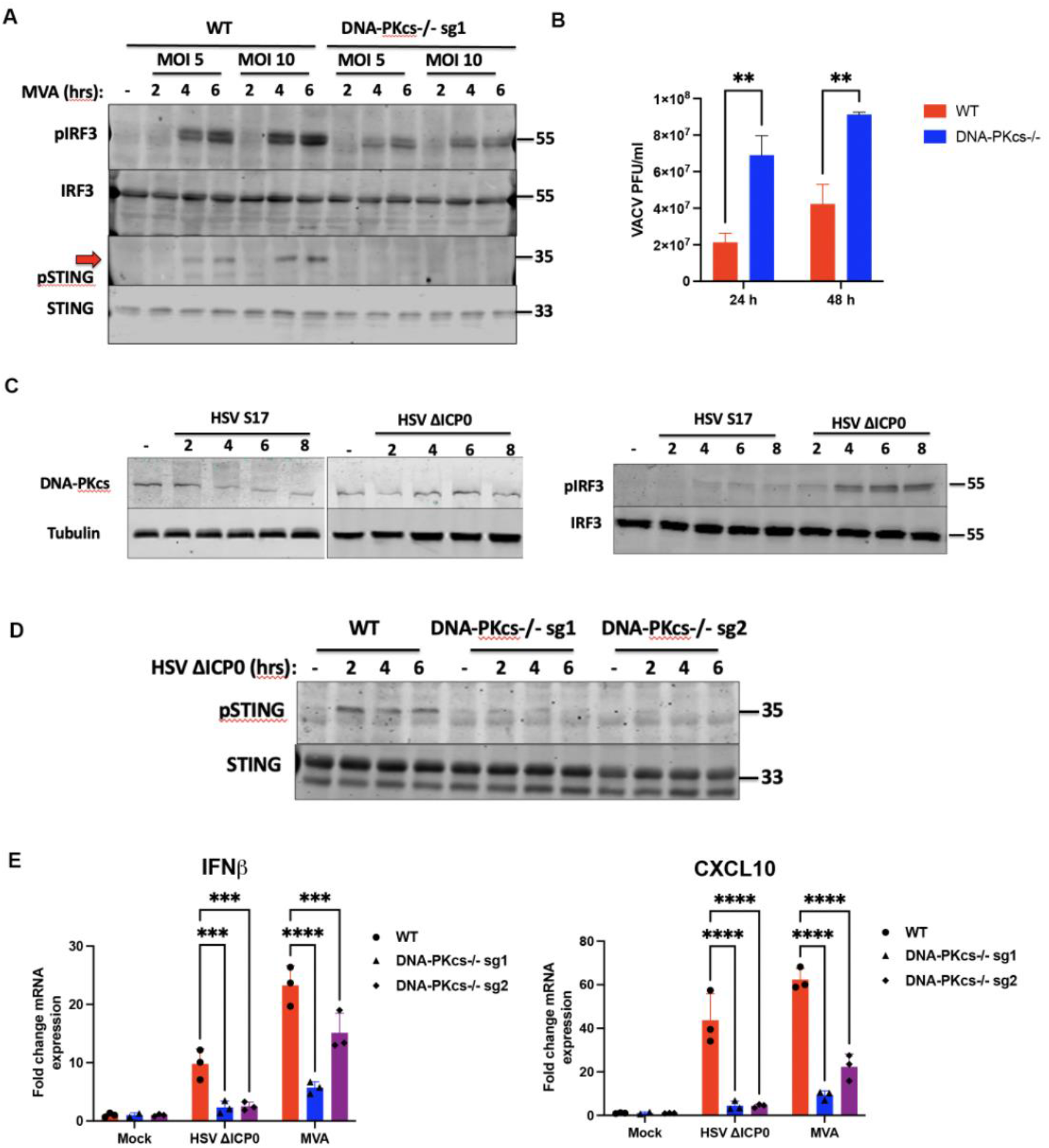
DNA-PKcs is required for innate sensing of DNA viruses in human cells. A) WT (DNA-PKcs^+/+^) and DNA-PKcs^−/-^ cells were infected with MVA at the indicated MOIs and analysed by immunoblotting with the indicated antibodies. Red arrow shows the position of the phospho-STING specific band. B) WT (DNA-PKcs^+/+^) and DNA-PKcs^−/-^ cells were infected with VACV TBio 6517 at MOI 0.01 and the production of infectious virions was quantified 24 or 48 h later by plaque assay on BSC-1 cells. C) HFFs were infected with WT (S17) or *dl1043* HSV-1 (ΔICP0) at MOI 5 and immunoblotted with the indicated antibodies. D) WT (DNA-PKcs^+/+^) and DNA-PKcs^−/-^ cells were infected with *dl1043* HSV-1 (ΔICP0) at MOI 5 and immunoblotted with the indicated antibodies). E) WT (DNA-PKcs^+/+^) and DNA-PKcs^−/-^ cells were infected with MVA or *dl1043* HSV-1 (ΔICP0) at the indicated times and analysed by qRT-PCR for *IFNB* and *CXCL10* 6 h post infection.

Next, we used herpes simplex virus 1 (HSV-1) strain 17, which we found not to activate IRF3 during HFF infection, but did degrade DNA-PKcs (Figure 6C), as previously reported^21^. We hypothesised that the degradation of DNA-PKcs may be interfering with the ability of the host cell to sense viral DNA and, therefore, we used the HSV-1 mutant *dl1043* virus that lacks expression of ICP0 (ref). *dl1043* infection of HFFs had no impact on DNA-PKcs expression but did result in measurable IRF3 activation (Figure 6C). Consistent with MVA infection, following *dl1043* infection wild-type HFFs could activate STING, but this was absent in DNA-PKcs KO cells (Figure 6D). In the absence of DNA-PKcs, MVA and *dl1043-*driven *IFNB* and *CXCL10* transcription were both significantly reduced compared with infection of wild-type cells (Figure 6E). In cells harbouring the L3062R mutation, *IFNB* and *CXCl10* transcription were enhanced in infected cells, consistent with this mutant having enhanced function in the context of STING-dependent signalling (Supplementary Figure 3). These data show that DNA-PKcs is required for the IFN-I response to *poxvirus* and *herpesvirus* infections consistent with its essential function in the ICD sensing pathway in human cells.

## Discussion

The function of DNA-PK in innate sensing of DNA and DNA viruses has been described in several contexts. In mice and in murine fibroblasts, DNA-PKcs, Ku70 and Ku80 are required for the intracellular DNA-driven IRF3 activation and for sensing vaccinia virus and HSV-1^4^. In monocytic THP-1 cells, Ku70 and Ku80 are required for sensing HSV-1 by triggering cGAS activation^8^. In HeLa cells, Ku70 interacts with the retrovirus human T cell leukaemia virus-1 (HTLV-1) reverse transcription intermediate DNAs to drive IRF3-dependent interferon beta responses^12^. In human endothelial cell lines, DNA-PK has been reported to regulate the interferon response to the herpesvirus KSHV^11^ and Ku70 is reported to regulate the chemokine response to hepatitis B virus (HBV) infection^30^. Our data here confirm the importance of DNA-PKcs in the sensing of both cytoplasmic and nuclear-replicating DNA viruses and its role in initiating the generation of a type-I interferon response to infection of human cells.

There is increasing evidence that the DNA-PK complex regulates one or more pathways downstream of ICD detection. In both murine and human fibroblasts, DNA-PKcs is required for STING-dependent IRF3 activation that leads to the generation of a classical anti-viral type-I interferon response. In this study, we show that human fibroblasts lacking DNA-PKcs fail to activate STING/IRF3 signalling and subsequent gene transcription in response to ICD stimulation and DNA virus infection. There are reports of a similar role of the Ku proteins in monocytic cells^8^ and of a potentially separate pathway that links Ku with IFN-III production, but that also requires STING^10,18^. In parallel, there is a report of DNA-PK activating a STING-independent pathway to IRF3 activation in a human leukaemia cell line^17^. The mechanisms underlying these co-operative or independent pathways are likely to involve direct interplay between DNA-PKcs/Ku70/Ku80 and other ICD sensing PRRs. Indeed, we and others^4,8,9,31^ have shown that Ku and/or DNA-PKcs can interact with cGAS and there is a reported interaction between DNA-PK and another ICD PRR, IFI16, that regulates the interferon response to HSV-1^7^.

Our data here, and in a previous report^4^, indicate that the kinase activity of DNA-PK is not required for its ability to activate the STING/TBK1/IRF3 signalling axis. Use of cells from SCID mice that have a deletion in the last 50 amino-acids of DNA-PKcs that inactivates the kinase domain^4^, or use of specific DNA-PKcs kinase inhibitors consistently show that there is no effect of blocking the kinase activity on ICD-driven signalling. As such, it is likely that the role of DNA-PKcs/Ku70/Ku80 in innate sensing is a structural one, possibly in sensing specific structures or sequences of DNA, such as DNA ends^17^ or nicks and gaps in DNA viral genomes in concert with IFI16^7,32^ and delivering them to cGAS to drive STING activation. In this context, it maybe that DNA-PK can positively or negatively regulate DNA sensing, depending on cell type and context of the infection^31^. Future studies will require more detailed structural and biochemical analyses of these processes and cell type-specific analyses using cells that are relevant to the infection processes being studied.

Further insight into the mechanisms by which DNA PRRs function in the context of infection can be obtained by studying the role of specific viral proteins that inhibit these host sensing pathways. During HSV-1 infection, DNA-PKcs is rapidly degraded^21^, although the Ku proteins remain stable, consistent with the concept that all three DNA-PK components are required for ICD sensing. Poxviruses encode the C16/C4 family of immunomodulatory proteins that use steric hindrance to block viral DNA from binding to the Ku proteins^19,20^, resulting in a reduction of the ICD-driven interferon response. This mechanism is consistent with a DNA-binding, structural function of DNA-PK in the context of intracellular DNA sensing. Notably, many other DNA viruses express proteins that bind or interfere with the function of DNA-PK^33^, although the function of many of these proteins in innate sensing is not yet described.

The discovery of patients with mutations in DNA-PKcs has provided further evidence for the function of this protein in innate sensing. The L3062R mutation analysed here causes severe combined immunodeficiency (SCID) by disrupting V(D)J recombination and lymphocyte maturation. Here, we show that primary fibroblasts from patients with this mutation hyperactivate STING following ICD stimulation, by an unknown mechanism. This mutation is on the surface of DNA-PKcs in the region of the protein that is responsible for binding Artemis^34^. It is hypothesised that this mutation disrupts DNA-PKcs/Artemis interactions and hence reduces the efficiency of double-strand DNA break repair and V(D)J combination. It is possible, but yet unexplored, that this DNA-PKcs/Artemis interaction may also regulate ICD sensing, in particular as Artemis has been implicated in manipulation of the terminal ends of viral genomes^35^. This contribution of DNA-PK-dependent DNA sensing to an auto-inflammatory phenotype is indicative of the requirement for exquisite balance in maintenance of homeostasis in relation to inflammatory signalling. The dysregulation of intracellular nucleic acid sensing contributes to multiple autoimmune and autoinflammatory diseases, such as Lupus and Aicardi-Goutieres syndrome^36,37^. It is possible that DNA-PKcs contributes to the pathogenesis of such disorders and, in a similar manner, to the triggering of anti-tumour immune responses^38^.

## Acknowledgements

We thank Professor Geoffrey Smith (Cambridge University, UK) for the MVA, Professor Gill Elliot (University of Surrey, UK) for the HSV-1 strain *dl1043* and Professor Mike Weekes (University of Cambridge, UK) for HFF-hTert cells. This work was funded by a Wellcome Trust Seed Award 201946/Z/16/Z and UKRI/BBSRC project grant BB/S001336/1 to BF.

## Author Contributions

D.H., E.W. and M.O. designed and performed experiments and analysed data. A.B. obtained patient samples. A.M., K.H. designed experiments, provided reagents, interpreted the data and edited the manuscript. B.F. coordinated the research, designed and conceived all experiments and wrote the manuscript.

## Supplementary Data

**Supplementary Figure 1.**
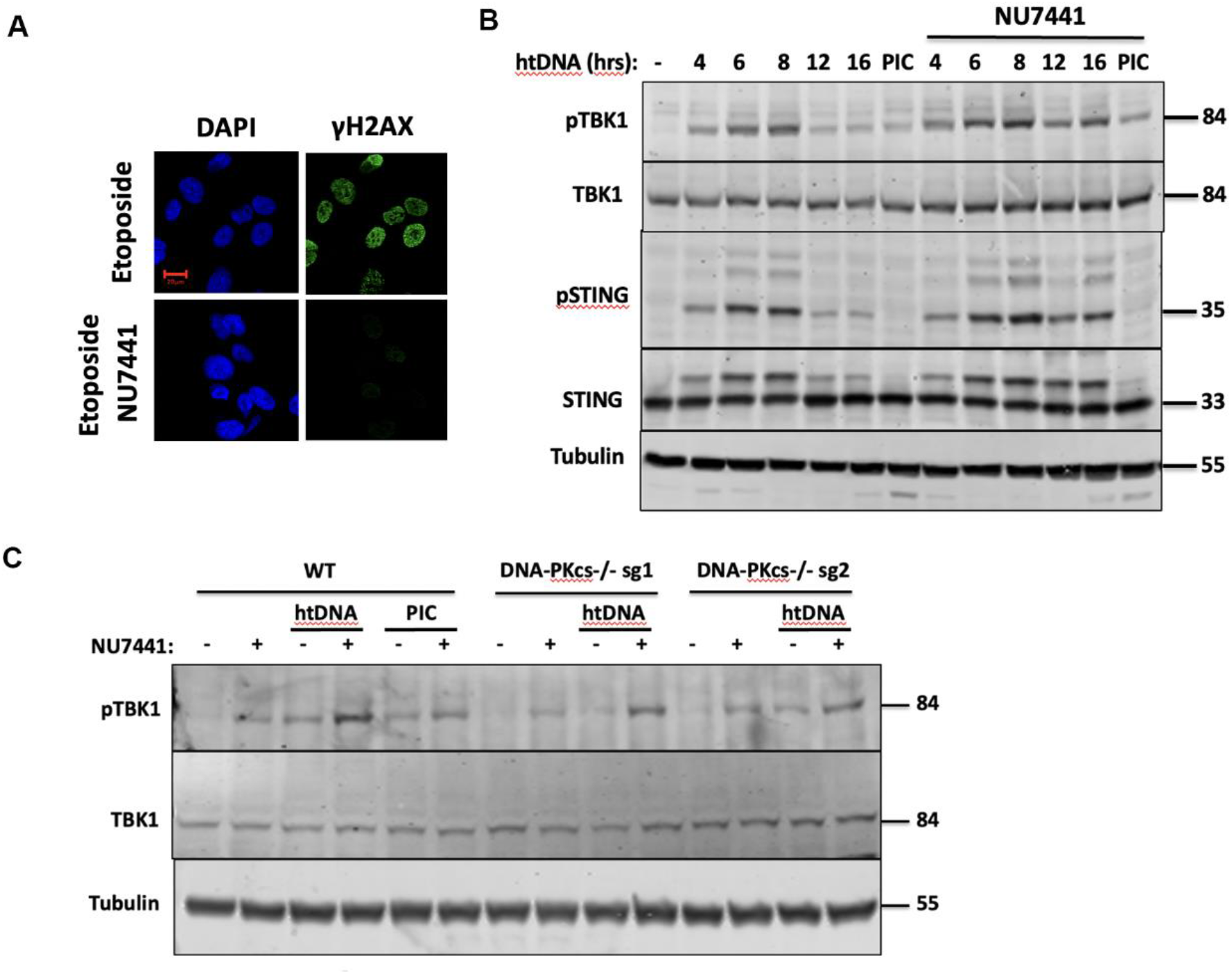
Nu7441 enhances ICD sensing by activating TBK1 independently of DNA-PKcs inhibition. A) HFFs were pre-treated with 2 μg/ml Nu7441 or carrier control and then with 30 μM etoposide for 2 h before being fixed and stained for anti-γH2AX. Scale bar=20 μm B) HFFs were pre-treated with 2 μg/ml Nu7441 or carrier control and stimulated with HT-DNA for the indicated times and immunoblotted with the indicated antibodies. C) WT (DNA-PKcs+/+) or DNA-PKcs −/− HFFs were pre-treated with 2 μg/ml Nu7441 or carrier control before stimulation with HT-DNA or poly(I:C) for 8 hours and then immunoblotted with the indicated antibodies.

**Supplementary Figure 2.**
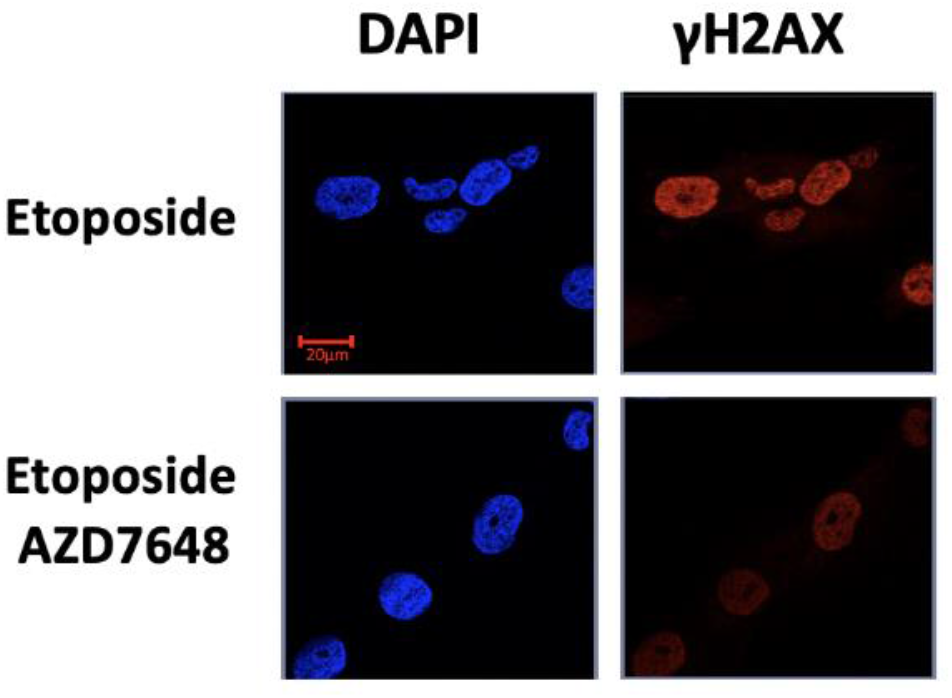
AZD7648 inhibits DNA-PKcs activity. HFFs were pre-treated with 2 μM AZD7648 or carrier control and then with 30 μM etoposide for 2 h before being fixed and stained for anti-γH2AX. Scale bar = 20 μm

**Supplementary Figure 3:**
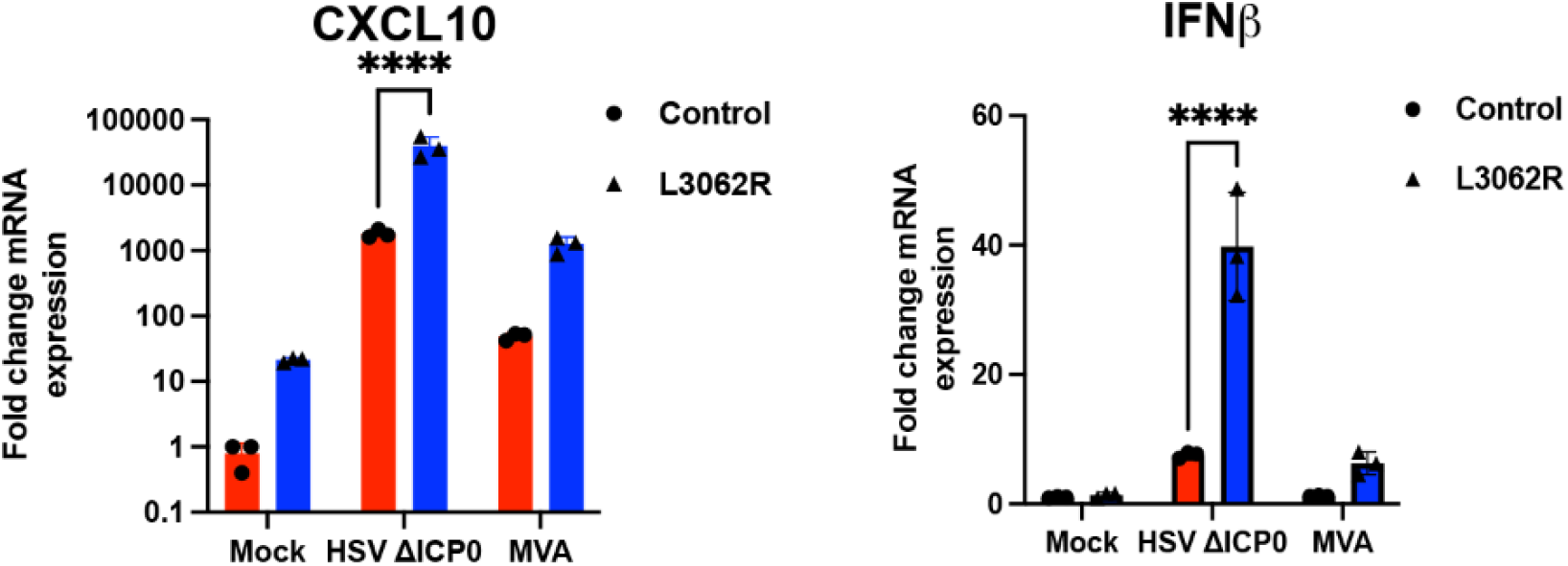
The human DNA-PKcs L3062 mutation enhances antiviral immune responses. Primary skin fibroblasts from healthy donor or a patient harbouring the L3062 mutation were infected with MVA or HSV-1 ΔICP0 and analysed by qRT-PCR for *IFNB* and *CXCL10* 6 h post stimulation.

## References

1. Mansur, D.S., Smith, G.L., and Ferguson, B.J. (2014). Intracellular sensing of viral DNA by the innate immune system. Microbes Infect. 16, 1002–1012. https://doi.org/10.1016/j.micinf.2014.09.010.

2. Iwasaki, A., and Medzhitov, R. (2015). Control of adaptive immunity by the innate immune system. Nat. Immunol. 16, 343–353. 10.1038/ni.3123.

3. Unterholzner, L., Keating, S.E., Baran, M., Horan, K.A., Jensen, S.B., Sharma, S., Sirois, C.M., Jin, T., Latz, E., Xiao, T.S., et al. (2010). IFI16 is an innate immune sensor for intracellular DNA. Nat. Immunol. 11, 997–1004. 10.1038/ni.1932.

4. Ferguson, B.J., Mansur, D.S., Peters, N.E., Ren, H., and Smith, G.L. (2012). DNA-PK is a DNA sensor for IRF-3-dependent innate immunity. Elife 1, e00047–e00047. 10.7554/eLife.00047.

5. Sun, L., Wu, J., Du, F., Chen, X., and Chen, Z.J. (2013). Cyclic GMP-AMP synthase is a cytosolic DNA sensor that activates the type I interferon pathway. Science 339, 786– 791. 10.1126/science.1232458.

6. Orzalli, M.H., Broekema, N.M., Diner, B.A., Hancks, D.C., Elde, N.C., Cristea, I.M., and Knipe, D.M. (2015). cGAS-mediated stabilization of IFI16 promotes innate signaling during herpes simplex virus infection. Proc. Natl. Acad. Sci. U. S. A. 112, E1773–81. 10.1073/pnas.1424637112.

7. Justice, J.L., Kennedy, M.A., Hutton, J.E., Liu, D., Song, B., Phelan, B., and Cristea, I.M. (2021). Systematic profiling of protein complex dynamics reveals DNA-PK phosphorylation of IFI16 en route to herpesvirus immunity. Sci. Adv. 7. 10.1126/SCIADV.ABG6680.

8. Tao, X., Song, J., Song, Y., Zhang, Y., Yang, J., Zhang, P., Zhang, D., Chen, D., and Sun, Q. (2022). Ku proteins promote DNA binding and condensation of cyclic GMP-AMP synthase. Cell Rep. 40, 111310. 10.1016/J.CELREP.2022.111310.

9. Patricio, D. de O., Dias, G.B.M., Granella, L.W., Trigg, B., Teague, H.C., Bittencourt, D., Báfica, A., Zanotto-Filho, A., Ferguson, B., and Mansur, D.S. (2022). DNA-PKcs restricts Zika virus spreading and is required for effective antiviral response. Front. Immunol. 0, 6130. 10.3389/FIMMU.2022.1042463.

10. Zhang, X., Brann, T.W., Zhou, M., Yang, J., Oguariri, R.M., Lidie, K.B., Imamichi, H., Huang, D.-W., Lempicki, R. a, Baseler, M.W., et al. (2011). Cutting edge: Ku70 is a novel cytosolic DNA sensor that induces type III rather than type I IFN. J. Immunol. 186, 4541–4545. 10.4049/jimmunol.1003389.

11. Morchikh, M., Cribier, A., Raffel, R., Amraoui, S., Cau, J., Severac, D., Dubois, E., Schwartz, O., Bennasser, Y., and Benkirane, M. (2017). HEXIM1 and NEAT1 Long Non-coding RNA Form a Multi-subunit Complex that Regulates DNA-Mediated Innate Immune Response. Mol. Cell 67, 387–399.e5. 10.1016/j.molcel.2017.06.020.

12. Wang, J., Kang, L., Song, D., Liu, L., Yang, S., Ma, L., Guo, Z., Ding, H., Wang, H., and Yang, B. (2017). Ku70 Senses HTLV-1 DNA and Modulates HTLV-1 Replication. J. Immunol., ji1700111. 10.4049/jimmunol.1700111.

13. Ishikawa, H., Ma, Z., and Barber, G.N. (2009). STING regulates intracellular DNA-mediated, type I interferon-dependent innate immunity. Nature 461, 788–792. 10.1038/nature08476.

14. Ablasser, A., Goldeck, M., Cavlar, T., Deimling, T., Witte, G., Röhl, I., Hopfner, K.-P., Ludwig, J., and Hornung, V. (2013). cGAS produces a 2’-5’-linked cyclic dinucleotide second messenger that activates STING. Nature 498, 380–384. 10.1038/nature12306.

15. KR, B., C, L., TL, S., AM, S., DJ, C., DB, D., F, M., M, T., KE, L., Y, Z., et al. (2020). TBK1 and IKKε Act Redundantly to Mediate STING-Induced NF-κB Responses in Myeloid Cells. Cell Rep. 31. 10.1016/J.CELREP.2020.03.056.

16. Fang, R., Wang, C., Jiang, Q., Lv, M., Gao, P., Yu, X., Mu, P., Zhang, R., Bi, S., Feng, J.-M., et al. (2017). NEMO–IKKβ Are Essential for IRF3 and NF-κB Activation in the cGAS– STING Pathway. J. Immunol. 199, 3222–3233. 10.4049/jimmunol.1700699.

17. Burleigh, K., Maltbaek, J.H., Cambier, S., Green, R., Gale, M., James, R.C., and Stetson, D.B. (2020). Human DNA-PK activates a STING-independent DNA sensing pathway. Sci. Immunol. 5. 10.1126/SCIIMMUNOL.ABA4219.

18. Sui, H., Zhou, M., Imamichi, H., Jiao, X., Sherman, B.T., Lane, H.C., and Imamichi, T. (2017). STING is an essential mediator of the Ku70-mediated production of IFN-λ1 in response to exogenous DNA. Sci. Signal. 10, eaah5054. 10.1126/scisignal.aah5054.

19. Peters, N.E., Ferguson, B.J., Mazzon, M., Fahy, A.S., Krysztofinska, E., Arribas-Bosacoma, R., Pearl, L.H., Ren, H., and Smith, G.L. (2013). A Mechanism for the Inhibition of DNA-PK-Mediated DNA Sensing by a Virus. PLoS Pathog. 9, e1003649. 10.1371/journal.ppat.1003649.

20. Scutts, S.R., Ember, S.W., Ren, H., Ye, C., Lovejoy, C.A., Mazzon, M., Veyer, D.L., Sumner, R.P., and Smith, G.L. (2018). DNA-PK Is Targeted by Multiple Vaccinia Virus Proteins to Inhibit DNA Sensing. Cell Rep. 25, 1953–1965.e4. 10.1016/J.CELREP.2018.10.034.

21. Parkinson, J., Lees-miller, S.P., and Everett, R.D. (1999). Herpes Simplex Virus Type 1 Immediate-Early Protein Vmw110 Induces the Proteasome-Dependent Degradation of the Catalytic Subunit of DNA-Dependent Protein Kinase Herpes Simplex Virus Type 1 Immediate-Early Protein Vmw110 Induces the Proteasome-Dependent De.

22. Stow, N.D., and Stow, E.C. (1986). Isolation and characterization of a herpes simplex virus type 1 mutant containing a deletion within the gene encoding the immediate early polypeptide Vmw110. J. Gen. Virol. 67, 2571–2585. 10.1099/0022-1317-67-12-2571/CITE/REFWORKS.

23. Davidson, S., Coles, M., Thomas, T., Kollias, G., Ludewig, B., Turley, S., Brenner, M., and Buckley, C.D. Fibroblasts as immune regulators in infection, inflammation and cancer. 10.1038/s41577-021-00540-z.

24. Correa-Gallegos, D., Jiang, D., and Rinkevich, Y. (2021). Fibroblasts as confederates of the immune system. Immunol. Rev. 302, 147–162. 10.1111/IMR.12972.

25. Fok, J.H.L., Ramos-Montoya, A., Vazquez-Chantada, M., Wijnhoven, P.W.G., Follia, V., James, N., Farrington, P.M., Karmokar, A., Willis, S.E., Cairns, J., et al. (2019). AZD7648 is a potent and selective DNA-PK inhibitor that enhances radiation, chemotherapy and olaparib activity. Nat. Commun. 2019 101 10, 1–15. 10.1038/s41467-019-12836-9.

26. Van Der Burg, M., IJspeert, H., Verkaik, N.S., Turul, T., Wiegant, W.W., Morotomi-Yano, K., Mari, P.O., Tezcan, I., Chen, D.J., Zdzienicka, M.Z., et al. (2009). A DNA-PKcs mutation in a radiosensitive T-B-SCID patient inhibits Artemis activation and nonhomologous end-joining. J. Clin. Invest. 119, 91–98. 10.1172/JCI37141.

27. Mathieu, A.L., Verronese, E., Rice, G.I., Fouyssac, F., Bertrand, Y., Picard, C., Chansel, M., Walter, J.E., Notarangelo, L.D., Butte, M.J., et al. (2015). PRKDC mutations associated with immunodeficiency, granuloma, and autoimmune regulator– dependent autoimmunity. J. Allergy Clin. Immunol. 135, 1578. 10.1016/J.JACI.2015.01.040.

28. Georgana, I., Sumner, R.P., Towers, G.J., and Maluquer de Motes, C. (2018). Virulent poxviruses inhibit DNA sensing by preventing STING activation. J. Virol. 92. 10.1128/JVI.02145-17.

29. Dai, P., Wang, W., Cao, H., Avogadri, F., Dai, L., Drexler, I., Joyce, J.A., Li, X.-D., Chen, Z., Merghoub, T., et al. (2014). Modified vaccinia virus Ankara triggers type I IFN production in murine conventional dendritic cells via a cGAS/STING-mediated cytosolic DNA-sensing pathway. PLoS Pathog. 10, e1003989. 10.1371/journal.ppat.1003989.

30. Li, Y., Wu, Y., Zheng, X., Cong, J., Liu, Y., Li, J., Sun, R., Tian, Z.G., and Wei, H.M. (2016). Cytoplasm-Translocated Ku70/80 Complex Sensing of HBV DNA Induces Hepatitis-Associated Chemokine Secretion. Front. Immunol. 7, 569. 10.3389/fimmu.2016.00569.

31. Sun, X., Liu, T., Zhao, J., Xia, H., Xie, J., Guo, Y., Zhong, L., Li, M., Yang, Q., Peng, C., et al. (2020). DNA-PK deficiency potentiates cGAS-mediated antiviral innate immunity. Nat. Commun. 11. 10.1038/S41467-020-19941-0.

32. Smith, S., Reuven, N., Mohni, K.N., Schumacher, A.J., and Weller, S.K. (2014). Structure of the herpes simplex virus 1 genome: manipulation of nicks and gaps can abrogate infectivity and alter the cellular DNA damage response. J. Virol. 88, 10146–10156. 10.1128/JVI.01723-14.

33. Turnell, a. S., and Grand, R.J. (2012). DNA viruses and the cellular DNA-damage response. J. Gen. Virol. 93, 2076–2097. 10.1099/vir.0.044412-0.

34. Niewolik, D., and Schwarz, K. (2022). Physical ARTEMIS:DNA-PKcs interaction is necessary for V(D)J recombination. Nucleic Acids Res. 50, 2096–2110. 10.1093/NAR/GKAC071.

35. Inagaki, K., Ma, C., Storm, T.A., Kay, M.A., and Nakai, H. (2007). The Role of DNA-PKcs and Artemis in Opening Viral DNA Hairpin Termini in Various Tissues in Mice. J. Virol. 81, 11304–11321. 10.1128/JVI.01225-07/ASSET/2BC9381A-A8F9-47B2-8C34-B5D417BEEFA3/ASSETS/GRAPHIC/ZJV0200797610009.JPEG.

36. Crow, Y.J., and Rehwinkel, J. (2009). Aicardi-Goutieres syndrome and related phenotypes: linking nucleic acid metabolism with autoimmunity. Hum. Mol. Genet. 18, R130–6. 10.1093/hmg/ddp293.

37. Ablasser, A., Hertrich, C., Wassermann, R., and Hornung, V. (2013). Nucleic acid driven sterile inflammation. Clin. Immunol. 147, 207–215. 10.1016/j.clim.2013.01.003.

38. Schadt, L., Sparano, C., Schweiger, N.A., Silina, K., Cecconi, V., Lucchiari, G., Yagita, H., Guggisberg, E., Saba, S., Nascakova, Z., et al. (2019). Cancer-Cell-Intrinsic cGAS Expression Mediates Tumor Immunogenicity. Cell Rep. 29, 1236–1248.e7. 10.1016/j.celrep.2019.09.065.

